# Automation of Experimental Workflows for High Throughput Robotic Cultivations

**DOI:** 10.1101/2023.12.05.570077

**Authors:** L. Kaspersetz, F. Schröder-Kleeberg, F. M. Mione, E. C. Martinez, P. Neubauer, M. N. Cruz-Bournazou

## Abstract

Process systems engineering methods and tools have been difficult to apply in bioprocess engineering, mainly due to the high complexity of biological systems and the low reproducibility of the experiments. High throughput robotic cultivation platforms in combination with computational tools for experimental design, resource scheduling, and operation, are rapidly gaining popularity. One important contribution being the generation of data in high throughput needed to overcome this lack of data with high information content and the worrying reproducibility crisis in life sciences. In this work, directed acyclic graphs are used to represent, manage and track all experimental workflows in a robotic platform. They support data provenance and enable traceability and reproducibility of workflows in robotic facilities. The experimental workflows are automated using Apache Airflow enabling to manage all necessary steps for fed-batch cultivations, including sampling, sample transport by a mobile robot, feed additions, data collection, storage in a SQL database and model fitting. The added value of this system is demonstrated in scale-down experiments, where *E. coli* BL21 (DE3), producing elastin like proteins, exhibits robustness towards glucose oscillations that mimic industrial cultivation conditions.

## 1 Introduction

The lack of high quality, reproducible experimental data in bioprocess engineering is still the bottle neck (Rogers et al., 2022) for scientific breakthroughs in bioengineering. To tackle this issue, robotic cultivation platforms optimally operated with computational tools can serve as generators of high-information content data. Such platforms rely on the integration of numerous interconnected devices, such as liquid handling stations, parallel cultivation systems and robots and their ability to generate the required data and provide provenance details regarding the rationale behind experimental design and the workflows used for conducting the experiments (Mitchell et al., 2022). A workflow management system (WMS) for automatically scheduling and executing the experimental and computational tasks not only increases the degree of automation but also contributes to implementing findable, accessible, interoperable and re-usable (FAIR) data principles (Wilkinson et al., 2016). Especially, in bioprocess development were scale-up decisions are based on small-scale experimental data, transparency and interoperability in experimental protocols and computational workflows can contribute to faster and less costly development throughout the different process scales. In this work, we present the implementation of a WMS based on directed acyclic graphs (DAGs) in a robotic cultivation facility.

## 2 Material and Methods

### 2.1 Bioxplorer cultivation platform

The BioXplorer 100 (H.E.L group, London, United Kingdom) composed of eight parallel cultivations in glass STRs, equipped with off-gas analyzers (BlueVary, BlueSens, Herten, Germany) was used for the cultivation workflows. The system is integrated into a liquid handling station (Tecan Group, Männedorf, Switzerland). A mobile robotic lab assistant (Astechproject Ltd., Runcorn, United Kingdom) consisting of a driving platform MIR100, a robotic arm URE5 equipped with a 2-finger gripper and 3D camera was used for automated sample transport. The reader is referred to Kaspersetz et al. (2022) for a detailed description of the platform and sampling procedures and at-line measurement procedures.

### 2.2 Strains

All experiments were carried out with *E. coli* BL21 (DE3) strains, carrying the pET28-NMBL-mEGFP-TEVrec-(V2Y)15-His expressing a recombinant fusion protein of an elastin like protein and eGFP, under the IPTG inducible lacUV5-promoter.

### 2.3 Cultivation

The preculture was set to an OD600 of 0.25 and cultured in 50 mL EnPresso B medium (Enpresso GmbH, Berlin, Germany) with 6 U L^−1^ Reagent A in 500 mL Ultra Yield flask sealed with an AirOtop enhanced flask seal overnight at 37°C and 220 rpm in an orbital shaker (25 mm amplitude, Adolf Kühner AG, Birsfelden, Switzerland). Main-cultures were run in four parallel glass stirred tank reactors (STR) each equipped with one Rushton type impeller at 37°C and pH was controlled at 7.0 with 7.5 % (NH_3*aq*_). The main cultures were started as 90 mL batch cultures with an initial glucose concentration of 5 g L-1. Two substrate addition strategies were used in the scale-down experiment: continuous feeding and bolus feeding profiles. The pulse-based feed followed a 10 min interval. The system was aerated with compressed air from 0.222 vvm to 1.666 vvm and stirring was increased from 1000 rpm 1450 rpm during the fed-batch phase.

### 2.4 Apache Airflow

Apache Airflow 2.2.4 and Python 3.7 was used to pro-grammatically describe, schedule and monitor the experimental cultivation workflows. The official Docker Image for Apache Airflow is hosted on DockerHub (apache/airflow:2.2.4). The workflows were authored as DAGs. All necessary for running an experimental workflow were included docker-compose.yml. All containerized applications were run via Docker Desktop 4.4.4 (73704) for Windows and a Windows Subsystem Linux 2 based back engine. The code is publicly available here.

### 2.5 *E. coli* growth model

The model is based on a mechanistic *E. coli* model with glucose partitioning, overflow metabolism, and acetate re-cycling. The mathematical model was formulated as an ODE system describing the changes in state variables for glucose, acetate, biomass, product and dissolved oxygen tension (DOT). The parameters of the model were obtained by fitting the model to the experimental data. A more detailed description of the underlying model and the functioning of the framework can be found in (Kim et al., 2023).

## 3 Device integration and worfklow automation

In bioprocess development, the implementation of automated cultivation workflows requires the integration of different devices, including mobile robots. Such robotic cultivation platforms usually consist of parallel bioreactor systems embedded in liquid handling stations (LHS) and analytical devices such as high-throughput analysers or additional LHS (Haby et al., 2019). The integration is often challenging due to missing standardized device interfaces or lack of adequate data infrastructure for handling the different data sources. Hence, managing and scheduling experiments in such facilities is a demanding task and inherently non-transparent without a workflow management system. In figure 1 a hierarchical infrastructure for the robotic cultivation platform is presented. All devices were integrated following a client-server architecture based on google Remote Procedure Call (gRPC) or Standardization in Lab Automation (SiLA2). The WMS in Apache Airflow (Harenslak & Ruiter, 2021) was implemented as the top layer. This digital infrastructure aims to provide a comprehensive tracking of the complete experimental workflow and to support transparency and reproducibility (Mitchell et al., 2022). All process data were stored in a SQL-database, while a shared file system was used to exchange data between containerized applications. The Apache Airflow web-service served as a user-interface for the experimental operator, allowing to trigger and monitor all experimental workflow steps.

**Figure 1:**
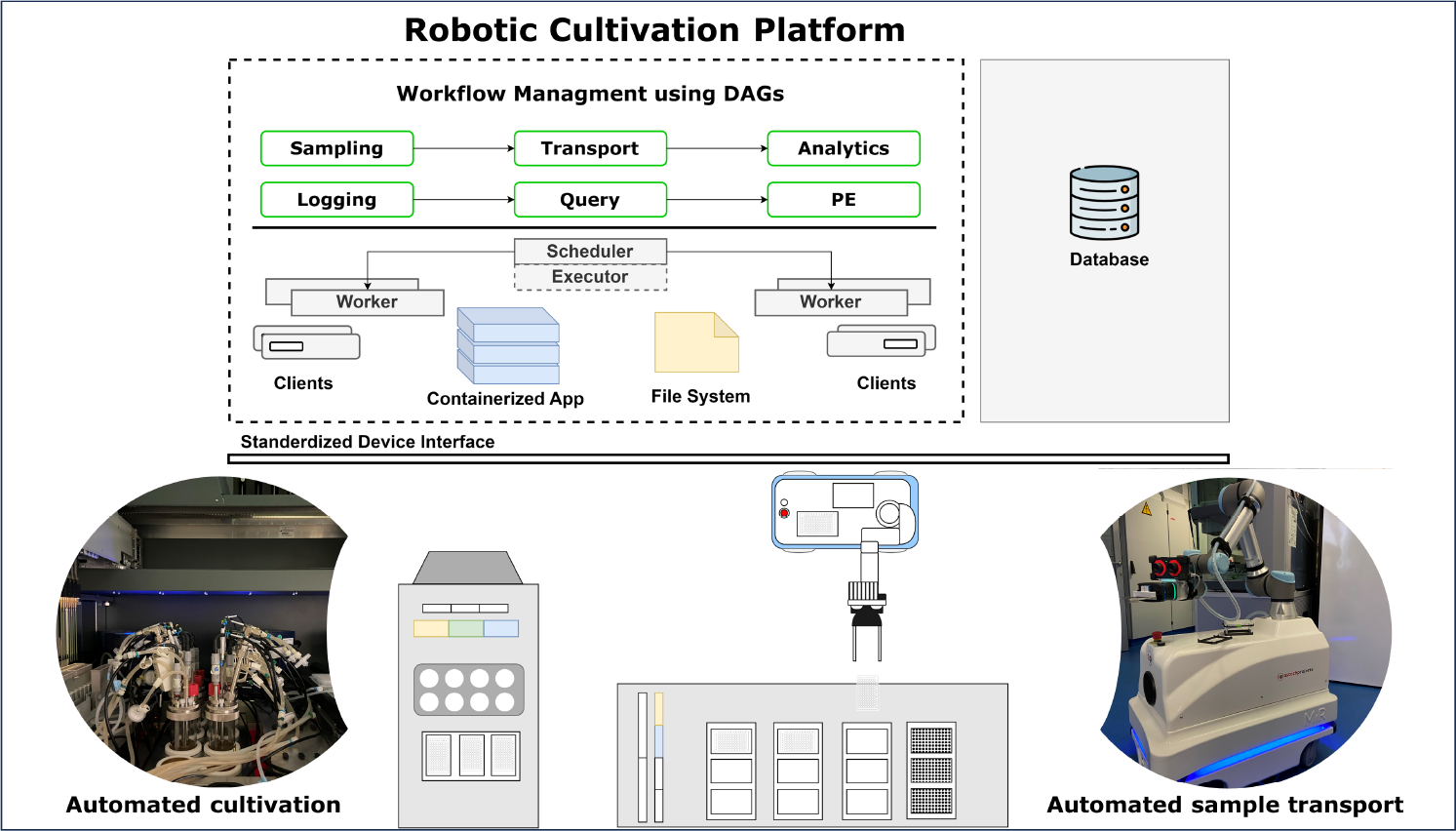
Schematic representation of the robotic cultivation platform with a workflow management system as the top layer.

**Figure 2:**
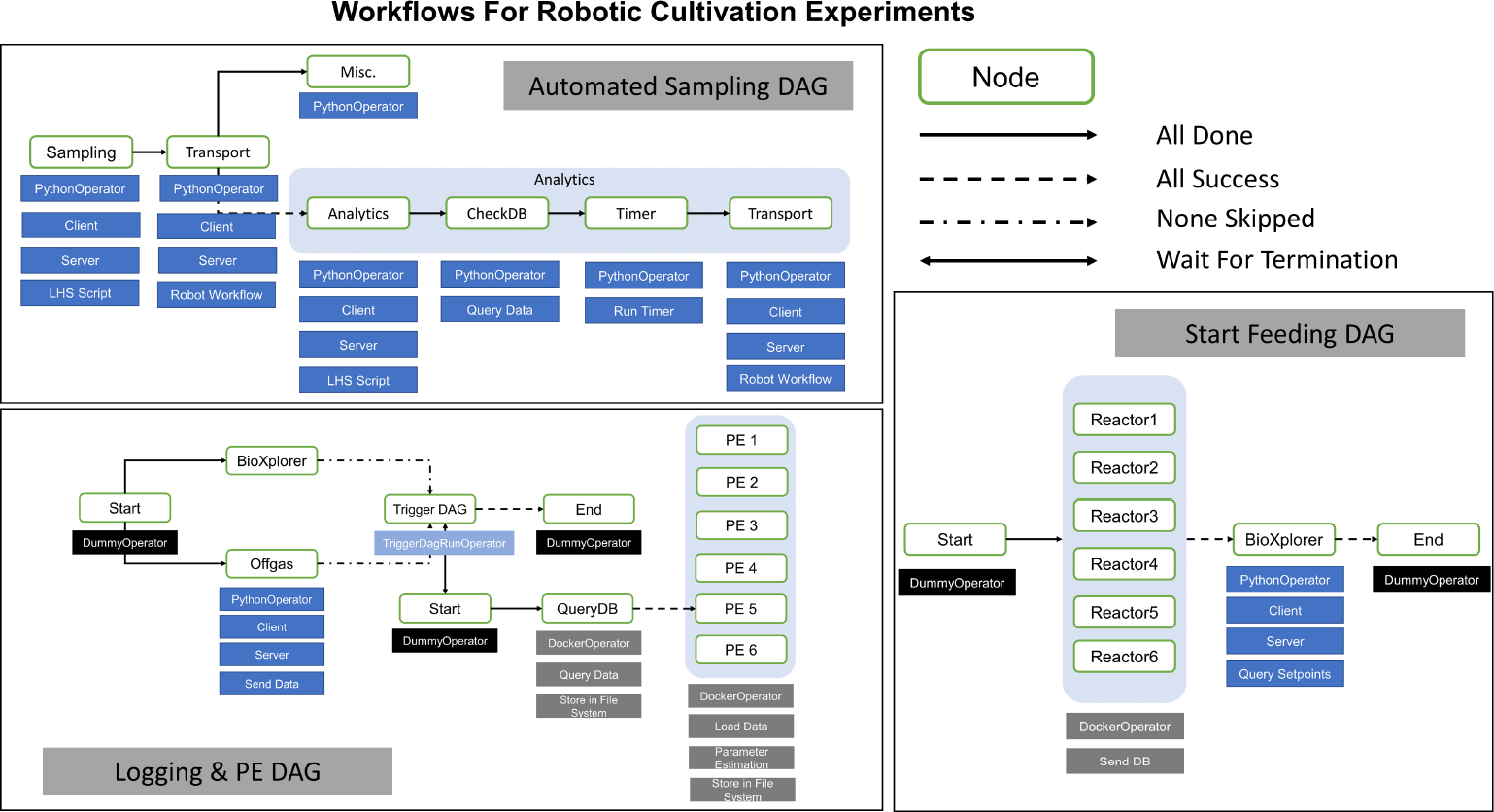
Representation of robotic cultivation workflows as directed acyclic graphs.

### 3.1 Implementation of experimental workflows

All Experimental workflows for managing fed-batch cultivations were represented through DAGs. The applicability of DAGs for enforcing FAIR principles by design in high throughput cultivation facilities has been recently addressed for computational workflows (Mione, Silva, Luna, Cruz B., & Martinez, 2022). In such a context, data provenance can be even increased if the complete experimental workflow starting from the device layer is incorporated into the WMS, allowing to understand how the experimental data was obtained and manipulated. The automated sampling DAG manages the sampling event of the LHS, the transportation of the microtiter plate by the mobile robot and the corresponding analytics. All nodes which are associated to a device, used the PythonOperator and the corresponding client for communication. The implementation allows for a six times loop over the whole sampling procedure. The DAG starts with a node that triggers the sampling procedure of the LHS. If the sampling event is successfully completed, the transportation node is carried out. After completion of the sample transport the corresponding analytics device is started, while in parallel the LHS can perform an additional task. Once the at-line analytics has been completed, the database is accessed to retrieve the corresponding results. The Logging and monitoring DAG manages the data needed by the cultivation system as well as queries data from the database for monitoring and updating the parameter estimation procedures. For the cultivation system and the off-gas analysis the implementation followed the same approach, using the PythonOperator and a client-server architecture. As soon as both devices are requested to send their online measurements to the database, the TriggerDagRunOperator starts the parameter estimation DAG. The computational pipeline for parameter estimation is executed inside a docker container. Containerized applications reduce integration effort, whilst increasing interoperability and reproducibility of computational workflows (Boettiger, 2015).

## 4 Case study – Scale down cultivation

Process performance parameters such as yields and titers of biotechnological process are often faltered when scaled up to industrial scale. This limitation occurs due to lack of robustness of the microbial host to perturbations in large-scale conditions (Olsson, Rugbjerg, Torello Pianale, & Trivellin, 2022). As cells move through the industrial-scale reactor, they are steadily exposed to changing environments. Hence, investigating the microbial response to such perturbations in usually homogenous and well-mixed small-scale reactors is necessary for a robust bioprocess development. Pulse-based scale-down bioreactor systems have shown to mimic gradients occurring in large-scale bioreactors to better characterize microbial responses and gather information about the strain robustness during scale-up. For details on scale-down simulators in high throughput systems, the reader is referred to Anane et al. (2019). The proposed infrastructure was used to manage the experimental workflows for conducting scale-down fed-batch cultivations to investigate the robustness of *E. coli* BL21 (DE3), producing elastin like proteins (Huber, Schreiber, Wild, Benz, & Schiller, 2014), to glucose oscillations. Two different feeding regimes were applied, following a continuous feeding or a pulse-based feeding regime. In figure 3 the cultivation data for parallel fed-batch cultivations with continuous feeding (R1 & R2) and pulse-based feeding (R3 & R4) is shown. The batch-phase ended after 5.3 h as indicated by glucose depletion and a rise in the O2-concentration in the off-gas. The exponential feeding was started using a µset = 0.2 h^−1^, leading to an oscillating pattern in the measured O2-concentration due to the pulse-based feeding. Throughout the feeding and induction phase, no glucose accumulation occurred. For the examined feeding strategies, no influence on the max. specific substrate uptake rate (qS_*max*_ = 1.5 g g h^−1^) and specific product (fig. 3) parameters were observed. However, in agreement with previous results (Anane et al., 2019) an increased maintenance coefficient (q_*m*_ = 0.09 g g h^−1^) for the pulse-based feeding strategy was observed.

**Figure 3:**
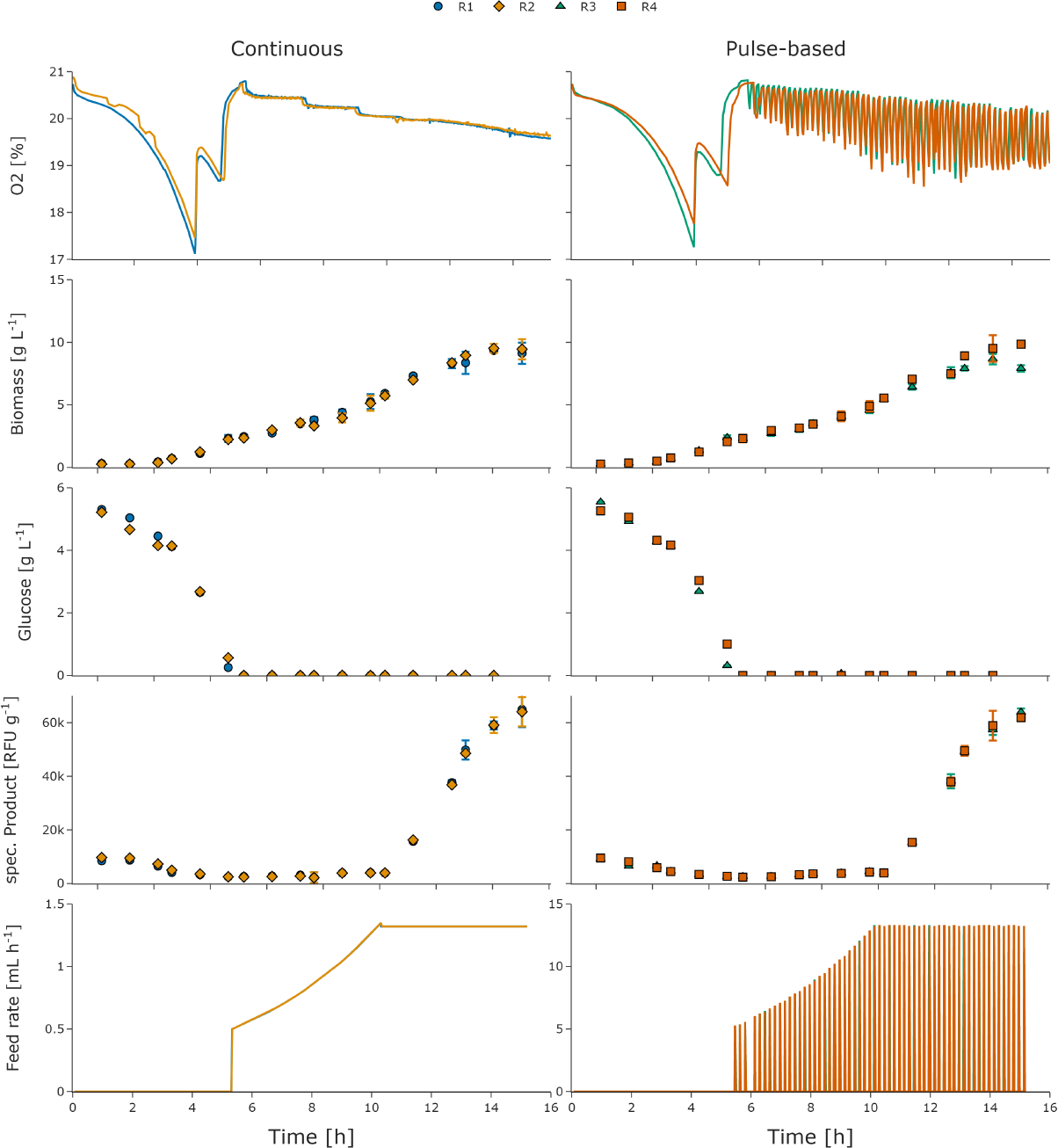
Parallel *E. coli* fed-batch cultivations, producing elastin like proteins, with continuous feeding (R1 & R2) in comparison to pulse-based feeding (R3 & R4).

## 5 Conclusions

We present the successful implementation of a WMS, based on DAGs, in a robotic cultivation facility. The proposed digital infrastructure is capable of scheduling and managing all necessary workflows for scale-down fed-batch cultivations, while increasing transparency and reproducibility of experiments and facilitating sharing of the conducted workflows. In the scale-down experiment, *E. coli* BL21 showed to be very robust with no adverse measurable physiological responses to glucose oscillations under the tested conditions, while fluorescence of product suggest a similar product quantity.

## 6 Acknowledgements

We gratefully acknowledge the financial support of the German Federal Ministry of Education and Research (01DD20002A – KIWI biolab).

